# FetchPA: a guided, end-to-end solution for local ATAC-Seq data processing and analyses

**DOI:** 10.64898/2026.09.10.750716

**Authors:** Dustin R. Fetch, Alexey A. Soshnev

## Abstract

Local genome accessibility strongly correlates with activity of cis-regulatory elements, and Assay for Transposase-Accessible Chromatin coupled with next-generation sequencing (ATAC-Seq) has emerged as method of choice to profile chromatin accessibility in both healthy and pathogenic conditions. The introduction of streamlined protocols and manufacturer kits has made this technique accessible to labs of a variety of disciplines, background, and research interests. Many bioinformatics tools have been created for the quality control, mapping, and visualization of ATAC-seq data, however these tools require familiarity with shell scripting, version control, UNIX directory structure, Python and/or R. Several pipelines for the processing of ATAC-seq data have been developed, yet even with these tools, bioinformatic analyses represents a bottleneck between wet-lab protocol execution and graphical representation of differentially accessible regions.

To address this problem, we assembled FetchPA, an intuitive pipeline which allows users with virtually no scripting and version control experience to install and manage all software for end-to-end analyses of ATAC-Seq data. FetchPA handles both local and public repository sources of sequencing data, executes standard QC benchmarks, and handles genome assembly and alignment using industry-standard PEPATAC pipeline. Further, it guides the user through the identification of differentially accessible regions and allows basic exploratory analyses via a dialogue interface. FetchPA operates in Windows Subsystem for Linux (WSL) and is installed via a single script that handles all individual tools, as well as their dependencies and updates, reference genome annotations and system resource allocation.

## 1. Introduction

Next-generation sequencing revolutionized the studies of gene regulation, and ATAC-Seq emerged as one of the most robust methods for quantitative characterization of regulatory elements at scale (Buenrostro et al., 2013; Buenrostro et al., 2015; Corces et al., 2017; Grandi et al., 2022). Robust and straightforward protocols have been developed for the wet-lab portion of the ATAC, including both commercially available kits and protocols to purify the key enzyme, Tn5 transposase, in-house (Soroczynski et al., 2024). Likewise, barcoded oligonucleotides compatible with Illumina sequencers for multiplex library preparations may be synthesized on-demand by commercial providers, and library purification and validation procedures are well established (Buenrostro et al., 2015). The sequencing process itself is typically outsourced to university core facilities or commercial service providers, and is equally robust. Yet in our anecdotal experience the bottleneck for ATAC-Seq implementation is downstream, and is defined by the relatively high barrier posed by bioinformatic analyses and data representation.

Despite intuitive workflow and many excellent pipelines available, *e*.*g*. nf-core/atacseq (Ewels et al., 2020; Patel et al., 2023), ENCODE (Hitz et al., 2023) and PEPATAC (Smith et al., 2021), even deployment of well-documented tools requires the end user to be comfortable with UNIX directory structure, command line, containers and version control – and thus represents a substantial barrier for users with limited experience in informatics. Alternative solutions include commercial software, often offered as a subscription service, or cloud-based tools which may offer limited analyses options, limited throughput, or be impractical if the original data must be analyzed locally due to privacy concerns.

To overcome these barriers, we previously developed a guided dialogue interface for RNA-Seq data analyses, FetchR (Fetch and Soshnev, 2026). Here, we describe a parallel toolkit for ATAC-Seq analyses based on the industry-standard <u>P</u>EP<u>A</u>TAC pipeline, FetchPA, aimed to lower the barrier for entry into ATAC-Seq analyses by the end user. FetchPA provides clear step-by-step guidance to all critical steps and offers straightforward analyses options, allowing for reasonably sophisticated exploratory analyses and data visualization. Last but not the least, it offers extensive documentation for reproducible outputs essential for data presentation and reporting.

## 2. FetchPA

Here, we briefly describe the FetchPA pipeline requirements, capabilities and output, and demonstrate representative analyses using a publicly available ATAC-Seq dataset.

### 2.1. System requirements and installation

The aim of developing FetchPA was to make every step of ATAC-Seq analysis available to end-user with limited to no bioinformatic experience. To this end, FetchPA facilitates easy installation of an industry-standard PEPATAC pipeline (Smith et al., 2021). Further, FetchPA allows for exploratory analyses to determine differentially accessible regions across experimental and control samples, and graphical representation of results for initial exploratory analyses driven by specific biological questions.

FetchPA operates under windows subsystem for Linux (WSL), accessible via Windows terminal in both Windows 10 and 11 operating systems. Installation of all relevant software is achieved through the execution of a single installer script. Prior to running, FetchPA installer ensures that sufficient disk space is available (estimated at a minimum of 10 GB), although this allocation would vary based on the amount of data end user intends to process. Installation begins with necessary system-wide packages (**Table 1**).

**Table 1:** System-wide packages installed by FetchPA.

| Software | Purpose |
| --- | --- |
| build-essential, git, pkg-config | Build toolchain |
| wget, curl, ca-certificates | Network/download utilities |
| bzip2, gzip | Compression tools |
| unzip | Archive/file utilities |
| perl | Scripting |
| libxml2-dev, libssl-dev, libcurl4-openssl-dev | XML/SSL/curl development |
| zlib1g-dev, libbz2-dev, liblzma-dev | Compression development |
| libuv1-dev | Asynchronous I/O library |
| fontconfig | Font configuration |
| coreutils, util-linux | Core System Utilities |

Following general system-wide installations, `Miniconda` is installed. Using Miniconda the installer creates a new `conda` environment, `pepatac`. All subsequent tools, with the exception of `Rust/gtars` and the `PEPATAC` repository, are installed within this environment (**Table 2**). Using Miniconda, the installer generates an environment named FetchPA within which command line tools and `R/python` packages are installed (**Tables 2** and **3**). `Rust/gtars` and the `PEPATAC` repository are installed separately.

**Figure 1.**
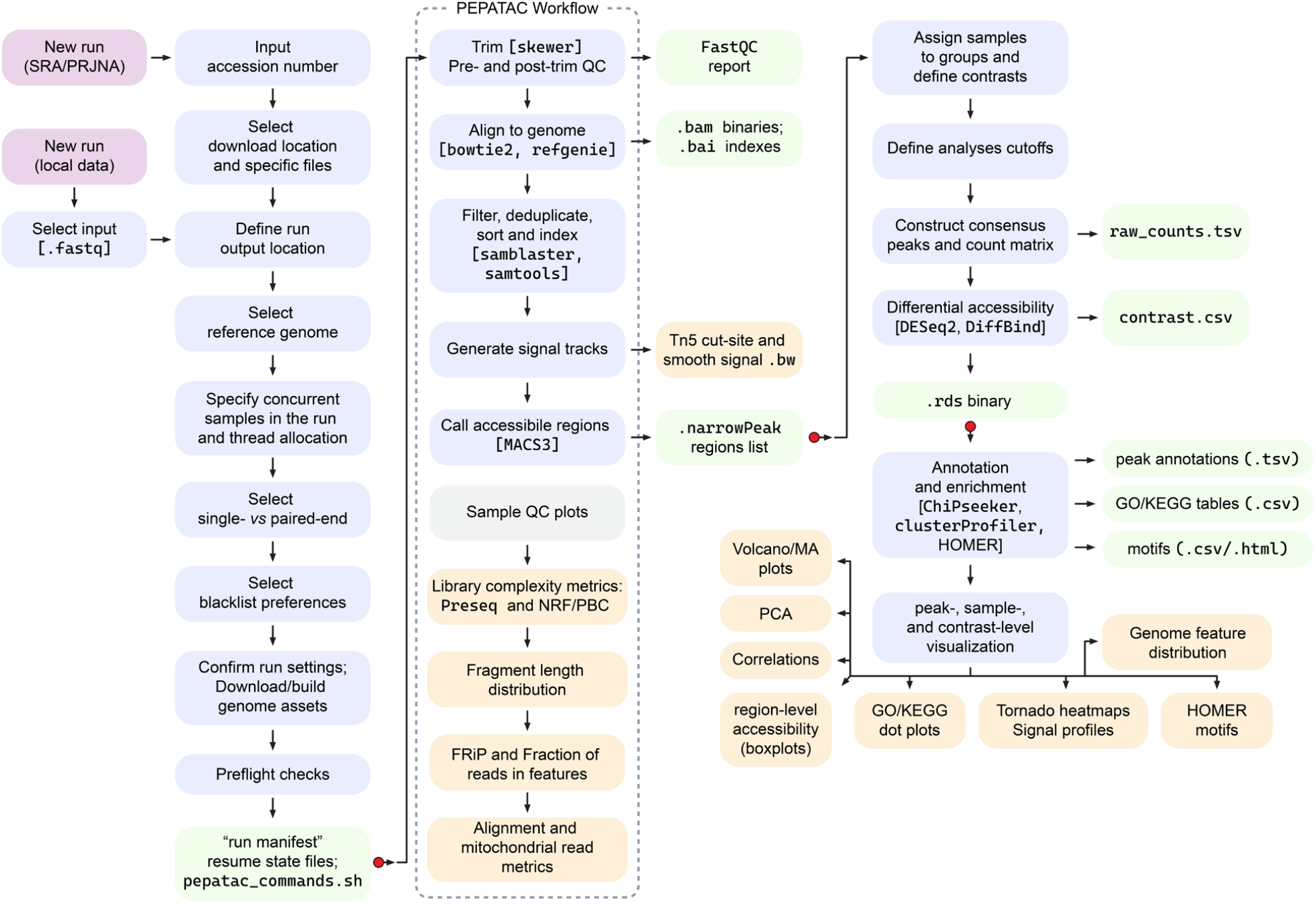
Workflow of FetchPA. FetchPA inputs include either local `.fastq` files or data from public repositories available via SRA/PRJNA accession numbers Input and settings are configured and saved in a “run manifest” and “sample manifest” files. `.fastq` files are processed with several consecutive QC checks, trimming, alignment and visualization via PEPATAC workflow. Regions list (.`narrowPeak`) output is analyzed via DESeq2 and subsequent exploratory analyses. Blue boxes indicate critical steps, green boxes denote significant outputs, and orange boxes show final visual outputs (see **Figure 2** for specific examples). Red dots indicate select “resume points” where analyses can be restarted without re-doing previously completed steps.

**Table 2:** Tools installed within FetchPA conda environment.

| Software | Pinned/default versions | Purpose |
| --- | --- | --- |
| bowtie2 | 2.5.4 | Short-read aligner |
| samtools | 1.21 | BAM filtering, sorting, indexing |
| bedtools | 2.31.1 | Genome arithmetic |
| fastqc | 0.12.1 | Pre/post-trim quality control |
| macs3 | 3.0.2 | Peak calling |
| deeptools | 3.5.6 | bamCoverage, computeMatrix, plotHeatmap |
| preseq | unpinned | Library complexity estimation |
| samblaster | 0.1.26 | Duplicate marking in stream |
| skewer | 0.2.2 | Adapter trimming (ATAC-specific) |
| trim-galore | 0.6.11 | Trimming wrapper |
| cutadapt | 5.2 | Trimming backend used by trim-galore |
| ucsc-wigtobigwig | unpinned | Wig to bigWig conversion |
| ucsc-bedtobigbed | unpinned | bed to bigBed conversion |
| homer | unpinned | Motif enrichment analysis |
| Refgenie | unpinned | Reference genome manager |
| Refgenconf | unpinned | Refgenie config library |
| pigz | unpinned | gzip-compatible compression/decompression |
| Wget, curl, unzip, tree, dos2unix | unpinned | General utilities |
| pip | 26.2 | Python package installer |
| r-base | 4.3 | R language |
| Python | 3.10 | Python language |
| Looper | 2.1.1 | PEPATAC pipeline execution |

At the start of the installation process the user is prompted to decide if they wish to continue using pinned software versions that are known to be compatible, or if they would prefer to check for more recent software versions. If the user opts to search for more recent software release, the pinned version of a given software is checked against current versions on conda-forge and bioconda channels.

If a newer version is available, the user is prompted with a choice to update or install the pinned version. Of note, this check only runs against software that have pinned versions, unpinned versions are left for `conda` to resolve freely to compatible versions. With `R` now installed within the `PEPATAC` environment, installation of `R`-specific packages follows (**Table 3**).

**Table 3:** Additional software and dependencies installed by FetchPA.

| Software | Purpose |
| --- | --- |
| r-remotes | Install r packages from Github/local |
| r-biocmanager | install Bioconductor packages |
| r-matrix, r-fs | Common utilities used by Bioconductor |
| r-xml, r-xml2 | XML parsing |
| r-bslib, r-rmarkdown | Report rendering |
| r-htmlwidgets | Interactive HTML output |
| r-shiny, r-dt | Interactive tables in PEPATAC reports |
| r-gplots | R plotting |
| r-pheatmap | Correlation and clustered heatmap generation |
| pepr | CRAN helper |
| optigrab | CRAN helper; commit 5166b8b |
| GenomicDistributions,<br>GenomicDistributionsData | Generates genomic distribution plots |
| DiffBind, edgeR, DESeq2, limma | Differential binding/accessibility analysis across sample groups |
| csaw | Required by DiffBind's background-bin normalization; version 1.36.0 |
| BiocParallel | Parallel computation framework used across Bioconductor packages |
| GenomicRanges | Manipulation of genomic intervals |
| Rsamtools | r interface to samtools/htslib for reading BAM files |
| SummarizedExperiment | Used for storing count matrices |
| AnnotationDbi | Used for accessing Bioconductor annotation databases |
| GenomicFeatures | Building/manipulating gene model feature sets |
| GenomeInfoDb | Genome/chromosome metadata |
| ChIPseeker | Peak annotation |
| ClusterProfiler, enrichplot | GO/KEGG enrichment |
| GO.db | GO term database backing clusterProfiler |
| `org.Hs.eg.db` | Genome-wide annotation database for human (Gene ID mapping) |
| `org.Mm.eg.db` | Genome-wide annotation database for mouse (Gene ID mapping) |
| `org.Rn.eg.db` | Genome-wide annotation database for rat (Gene ID mapping) |
| `org.Dm.eg.db` | Genome-wide annotation database for fruit fly (Gene ID mapping) |
| `org.Dr.eg.db` | Genome-wide annotation database for zebrafish (Gene ID mapping) |
| `TxDb.Hsapiens.UCSC.hg38.knownGene` | Human gene models (hg38) |
| `TxDb.Mmusculus.UCSC.mm10.knownGene` | Mouse gene models (mm10) |
| `TxDb.Rnorvegicus.UCSC.rn7.refGene` | Rat gene models (rn7) |
| `TxDb.Dmelanogaster.UCSC.dm6.ensGene` | Fruit fly gene models (dm6) |
| `TxDb.Drerio.UCSC.danRer11.refGene` | Zebrafish gene models (danRer11) |
| `ggrepel`, `dplyr`, `tidyr` | General data handling/visualization |
| `rust` (`rustc/cargo`) | Toolchain required to build gtars from source |
| `Gtars` | Fragmentation scoring / uniwig; git tag v0.9.0 |
| `IRanges` | Genomic interval representation/manipulation |
| `RSQLite` | SQLite database interface required by annotation infrastructure |
| `PEPATACr` | R reporting package accompanying PEPATAC; commit: 69c3fad7226152a5f4e040657cd1cb22a4aef6c5 |
| `Libxml2`, `pkg-config`, `libuv` | C libraries |

The script then clones, or if present already, retrieves the `PEPATAC` repository (commit 69c3fad) as well as all Python packages necessary to run PEPATAC later. The `PEPATACr` R package is taken directly from the cloned repository. Finally, `Refgenie` is initialized. With all installations completed the installation script writes another small script, `pepatac_check.sh`. This script optionally checks that all command line interface tools and `R` packages are present, and verifies the expected `csaw` and `gtars` version, as well as that `Refgenie` is present and accurately configured. The installer itself runs an equivalent check before reporting that all installation has been successful. To help facilitate reproducibility and ease of installation to other machines, both a minimal and full environment are exported following installation.

### 2.2. Data import and processing

With installation complete, the FetchPA pipeline now utilizes a second runner script which is responsible for the execution of the industry standard PEPATAC pipeline. PEPATAC performs read preprocessing using `skewer`, then aligns trimmed reads to a user-indicated reference genome using `Bowtie2` (Langmead and Salzberg, 2012), removes mitochondrial reads, and calls peaks using `MACS3` (Zhang et al., 2008). FetchPA explicitly provides PEPATAC with an assembly-appropriate effective genome size for peak calling instead of defaulting to the effective human-genome size. For genomes with an available standard blacklist (hg38, dm6, and mm10), users can choose to enable blacklist filtering, wherein the blacklist regions are automatically downloaded, and peaks overlapping with blacklisted regions are removed. For genomes lacking these standardized lists users may provide a custom blacklist. Among other outputs, the runner provides a .`narrowPeak` file, `.bam/.bai` files, and `.bigWig` file for each provided sample. The runner script then writes a QC summary file which details the finished run, as well as a sample sheet formatted to be used by the downstream differential analysis script. A sample manifest, written at the start of every run, details what samples were processed and in which way they were processed for future reference, ensuring reproducibility of reporting. A separate machine-readable resume state file allows for resumption of interrupted runs, restarting processing with previous user indicated configurations.

The runner accepts`.fastq` files stored locally on the machine or those downloaded using an SRA study (SRP) or BioProject accession number. Files retrieved this way are downloaded to a user-specified location and checksum validated using an MD5 checksum. Importantly, users are not required to download all sequencing files associated with an SRP/PRJNA accession number, instead only downloading and processing identified files of interest.

After indicating which files to process, user would indicate the matching organism. This prompts FetchPA to select reference genome assembly to use during data processing. For hg38, mm10, and dm6 genomes users may opt to download indexed genome assembly directly. Other genomes, including rn7 and danRer11, are currrently lacking this option - however the runner script would download the reference genome from UCSC portal and create an indexed reference using `Bowtie2`. Users may also use this option to generate a local index for hg38, mm10, and dm6 genomes should they wish. Users may also use other UCSC assemblies by providing a valid assembly identifier along with assembly-matched TxDb and species-matched OrgDb packages. These resources are validated and stored by the runner before processing begins.

**Figure 2.**
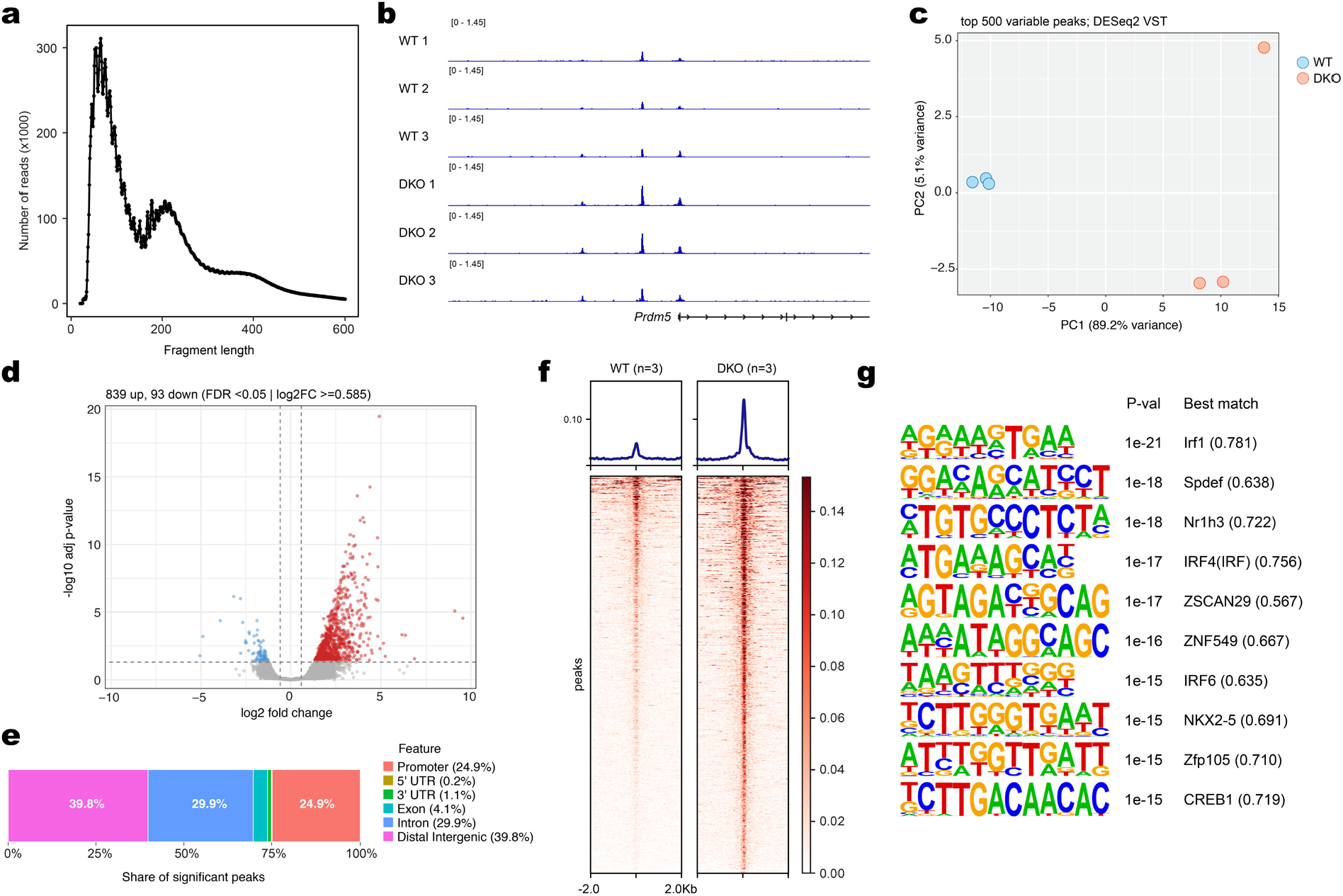
Representative outputs of FetchPA. **a**, fragment size distribution plot from a single ATAC-Seq library; **b**, .bigwig visualization of individual replicates; **c**, principal component analyses plot for all samples in the dataset **d**, volcano plot for visualization of differentially accessible regions under user-defined parameters and contrasts, **e**, distribution of differentially accessible regions relative to genome features, **f**, tornado plots of signal at differentially accessible regionsб, showing median signal of three replicates per condition; **g**, motifs enriched at differentially accessible regions. All examples shown here represent direct output of FetchPA minimally adjusted in Adobe Illustrator, using ATAC-Seq data from (Yusufova et al., 2021), downloaded from Gene Expression Omnibus GSE143293.

Because `PEPATAC` processes samples independently, FetchPA allows for multiple `PEPATAC` instances to be launched concurrently. The user chooses how many instances to launch as well as how many threads to allocate to each `PEPATAC` process, processing multiple samples in parallel. Before the run begins, users are prompted with a final display of samples to be processed and an overall run configuration summary. With the user’s confirmation, the runner begins processing the raw data.

### 2.3. Differential peak analyses

FetchPA differential analysis begins by reading a sample sheet written by the runner script, `samples_for_R_autodetected.csv`. This contains a list of all samples and where the differential script can locate them. After users provide the sample sheet to the `diff_analysis` script, they are offered an option to exclude samples from analysis. This is important, as all included samples are used to create a shared consensus peak set and read-count matrix. All samples included at this step will also contribute to the global PCA generated from the shared count matrix. With that in mind, only relevant samples should be included in a given differential analysis script run. Users then assign replicate samples to groups, and define one or more pairwise contrasts between previously named groups. Importantly, the differential analysis script requires at least two samples per group, those contrasts containing fewer than two samples in a given group are automatically skipped. Reference genome assembly is confirmed, with the genome used for alignment being autodetected and provided as a default. The script then confirms that it has access to both the TxDb and OrgDb for the selected genome, using the frozen snapshots associated with the particular run in question when available. The user then chooses whether or not to perform motif analysis on detected significantly differentially accessible regions using HOMER (Heinz et al., 2010). In this setting, users can opt to only run HOMER against known motif signatures, perform *de novo* motif discovery, or both. Of note, HOMER is configured to run on a combined significant peak list, as well as separate sets of peaks that exhibit increased and decreased accessibility.

The user then sets analysis parameters. The first, minimum peak overlap, determines the minimum number of samples a peak must appear in to be included in the consensus peak set. The `DiffBind` recommended value of 2 is provided to users as a default which can be adjusted at user discretion for more or less stringent definitions. User also defines statistical cutoffs provided as FDR-adjusted p-value, and log2-transformed fold change required to call significance. Users then determine resource (processor thread) allocation, as well as where they wish to save results. A final run summary is provided where all run information is displayed. With the user’s approval, differential analysis begins. Although a shared consensus peak set is constructed across all included samples, each pairwise contrast is performed on a subset of the two groups being compared. This means that samples are independently normalized and analyzed separately of all other samples. Although not dictated or influenced by user input, significant peaks are also annotated using ChIPseeker (Yu et al., 2015), linking peaks to genomic features and nearby genes. GO and KEGG are also both run on genes associated with significant peaks, including those of increased accessibility, decreased accessibility, and a combined set. Importantly, although this script does the majority of computational work necessary for data visualization, figure production is left to the subsequent explorer script, which requires the `explorer_bundle.rds` input file written at the end of every differential analyses run.

### 2.4. Exploratory analyses

The explorer script allows users to visualize their data, creating and recreating plots in a fashion that requires minimal system resources from their computer. The only exception is in the generation tornado plots, discussed below, which utilize `deepTools` and benefit from increased resource allocation. With nearly all computational heavy lifting complete, the explorer `.rds` file is portable, allowing for rapid visualization and investigation of the user’s data on nearly any PC. The `.rds` file contains the full-dataset VST matrix, sample metadata, completed contrast information, differential-accessibility results, analysis thresholds, as well as paths to the associated differential-analysis output and sample `.bam` files. Other results, including peak annotations, GO/KEGG enrichment results, and motif analysis results remain in the differential analysis output directory. The explorer script accesses these files as necessary during analysis and visualization. In total, the explorer script is capable of generating: PCA plots and correlation heatmaps using the contained VST matrix; volcano/MA plots from the per-contrast differential accessibility results saved in the explorer bundle; peak accessibility boxplots using the raw consensus-peak count matrix and associated normalization information saved separately from the explorer bundle in the differential-analysis output directory; HOMER motif summaries, GO/KEGG summaries, annotation distribution plots, and tornado plots generated from files in the differential-analysis output folder. All plots are customizable by the user, including sample selection for PCA and sample correlation analysis, customize FDR/log2 thresholds in volcano and MA plotting, select specific genomic coordinates for peak boxplot generation, and select genomic reference points when making tornado plots. The explorer script, as a whole, is meant to expedite initial analysis of data, streamlining exploration and discovery.

## 3. Concluding remarks

FetchPA was designed to make ATAC-Seq data processing and analyses accessible to a user with limited experience in bioinformatics. This pipeline is not intended to substitute the knowledge of statistics and informatics as these are imperative for the proper handling of data and interpretation of results. Instead, we understand that for new users, there exists a knowledge and experience gap when first performing genome-wide analysis, which we envision this pipeline can help bridge.

## Acknowledgements

We thank the members of the Soshnev laboratory at UT San Antonio for helpful suggestions and testing of FetchPA. The code was written with the assistance of Claude AI. The authors take full responsibility for the code and its annotation.

## Author contributions

D.R.F. conceived the project and designed FetchPA with input from A.A.S; D.R.F and A.A.S jointly wrote the manuscript.

## Data and code availability

FetchPA is available at https://github.com/DustinFetch/FetchPA-ATACseq-Pipeline

